# Plant diversity shapes plant volatile emission differently at the species and community level

**DOI:** 10.1101/2025.04.30.651392

**Authors:** Pamela Medina-van Berkum, Cynthia Albracht, Maximilien Bröcher, Marcel Dominik Solbach, Gideon Stein, Michael Bonkowski, François Buscot, Anna Heintz-Buschart, Anne Ebeling, Nico Eisenhauer, Tarek S. El-Madany, Yuanyuan Huang, Karl Kuebler, Sebastian T. Meyer, Jonathan Gershenzon, Sybille B. Unsicker

## Abstract

Studies have investigated the interactions between plants through competition and resource sharing to understand the mechanisms behind the positive effects of plant diversity on productivity. Volatile organic compounds (VOCs) are important info-chemicals in plant-plant interactions, but they have so far rarely been considered in this context. Here, we measured VOC emissions at both community and species levels (*Plantago lanceolata*) in experimental plant communities of varying diversity (The Jena Experiment) to understand the role of VOCs in driving biodiversity-ecosystem functioning relationships. We show that plant diversity determines the release of plant VOCs at both levels. At the community level, plant species richness directly enhanced VOC emission and increased VOC richness both directly and indirectly by altering herbivore damage and LAI. At the species level, plant diversity did not directly affect the VOC emissions of *P. lanceolata* but indirectly affected it by influencing the VOC emissions from the surrounding community. *Plantago lanceolata* individuals in communities with high concentrations of green leaf volatiles decreased their VOC emission, while those in communities with high concentrations of terpenoids increased their VOC diversity. Our results provide first evidence that plant diversity shapes community-level plant VOC emission and thus influences focal plant VOC emission inside the community.

## Introduction

Understanding how plant diversity affects community structure and interactions has been a major driver of research in biodiversity and ecosystem functioning relationships in recent years. Field experiments have shown that plant diversity not only increases plant community productivity but also causes metabolic changes in both primary and specialized metabolites in grassland plant species^1–3^. As a result, individuals of the same species growing in environments with varying diversity may exhibit differences in their trait expression, such as in their metabolite profiles, as they experience differing biotic pressures^4–6^.

Plants emit a large variety of volatile organic compounds (VOCs), including green leaf volatiles, terpenoids and aromatics^7^. These metabolites are released by all types of plant organs in response to both biotic and abiotic stimuli^8^ and they fulfill a plethora of functions. For instance, VOCs can act as signals for pollinators, parasitoids, or mycorrhizal fungi, as defense compounds to repel antagonists like herbivorous insects and pathogens^9–11^ and as important signals in intra- and inter-specific plant communication^12,13^. By mediating these interactions, VOCs might enhance the plant adaptability to a variety of environmental challenges^14^ and thus provide indicators of changes in their abiotic and biotic environment.

The variety of VOCs arises from the participation of several different core pathways and possibly additional enzymes^7^. VOC diversity can thus vary in terms of the number of compounds, their relative abundance, and their biosynthetic origins. Although plants emit VOCs constitutively, the high diversity of their emitted blends also arises from responses to other organisms and abiotic environmental factors^14^. Numerous studies have shown that herbivore or pathogen attacks drastically change plant VOC profiles quantitatively and qualitatively^15–17^. Moreover, environmental factors, such as drought and light limitation, have also been recognized to influence the VOC profiles of the plants^18^. Consequently, the complexity of the surrounding plant community, moderated by plant diversity, might shape their VOC emissions.

With increasing plant diversity, herbivore-inflicted leaf damage increases^19^, while pathogen infection is reduced^20,21^. This raises the question of what role VOC emissions play in this context. While most VOC studies have focused on the plant species level^5,22,^ the effects of plant diversity on VOCs emission can occur at different scales, with factors influencing patterns at the species level differing from those at the community level. At the species level, leaf damage can directly influence VOC profiles, but it can also have indirect effects. Exposure to herbivore- or pathogen-induced plant volatiles, for example, can trigger changes in the VOC profiles of nearby receiving plants and consequently reduce subsequent herbivore damage^23–25^. Notably, these responses can vary depending on the neighboring community and their identity^22,26^. Previous research found that *Trifolium pratense* L. altered its VOC diversity in response to both intra- and interspecific competition, suggesting that community diversity influences VOC-mediated herbivory responses^23^. Similarly, *Plantago lanceolata* L. showed a decreased VOC diversity with increasing plant diversity^27^, further supporting the role of community composition in shaping VOC profiles.

At the plant community level, research on VOC emission has largely been theoretical, with limited experimental evidence. Plant diversity could directly influence VOC complexity through the increase of plant species richness or plant phylogenetic diversity^27^. However, it may also have indirect effects on VOC profiles by modifying below- and above-ground abiotic and biotic factors. For example, increasing plant species diversity can increase herbivore load and reduce soil pathogens^19,28,^ and both herbivores and soil pathogens are known to influence plant VOC emissions^14^. Additionally, increased species diversity can affect abiotic conditions such as light availability, soil temperature, and moisture^29^. An increase in leaf area index (LAI) or vegetation height can reduce light availability, thereby affecting the emission of light-dependent compounds such as terpenoids^30^. These hypotheses offer alternatives to how increasing plant diversity could directly and indirectly influence VOC emissions at the community level, which ultimately may affect emissions at the species level as well.

Despite their ecological importance, most of the research on VOCs has been limited to controlled environments in laboratories or greenhouses. Experts have long called for more studies under natural field conditions^14,27,31,^ but only a limited number of research projects have explored the VOC profiles of plants under natural ecosystems^31–33^. Furthermore, most of these studies have focused on individual plants and species, with few examples at the community level^34^. Studies of VOC emission by communities with varying numbers of species are especially lacking. Since changes in VOC profiles, both at the species and community levels, may have a major influence on ecosystem functioning by altering interactions with other organisms, it is important to better understand the factors affecting VOC emission under field conditions in complex plant communities.

In this study, conducted in the framework of a long-term grassland biodiversity experiment (The Jena Experiment^35^), we investigated how VOC profiles vary across an experimental plant diversity gradient at both the community and the species levels, focusing specifically on *Plantago lanceolata* L. We selected this species due to the available insights into its metabolomic profiles, ecological interactions with other organisms, responses to plant diversity, and sufficient replication across the plant diversity gradient^5,36,37^. Here we show that community-level VOC diversity increases with increasing plant diversity, whereas at the species-level, plant diversity does not directly influence the VOC emissions of *P. lanceolata* but rather affects them indirectly by shaping the VOC emissions from the surrounding community. Our results provide insight into the various factors shaping VOC diversity across a plant diversity gradient, highlighting the distinctions between species- and community-level VOC patterns.

## Results

### Community VOC emission and richness increase with increasing plant species richness

First, we investigated whether the initially sown gradient in plant species richness, which ranges from monocultures to 60-species mixtures, is still present in the communities under study. In total, 82 plant species belonging to 61 genera, 19 families, and 14 orders were identified in the communities. Twenty-three of these species colonized the communities from the regional species (not belonging to the original species pool of the Jena Experiment, Table S1). Realized plant species richness (Hill q0), taxonomic diversity (Hill q1) and phylogenetic diversity (Hill q0 and q1) in the communities were positively correlated with the originally sown plant richness (taxonomic richness: R^2^ = 0.45, *p =* 0.011; taxonomic Shannon diversity: R^2^ = 0.30, *p =* 0.013; phylogenetic richness: R^2^ = 0.35, *p =* 0.005, phylogenetic Shannon diversity: R^2^ = 0.35, *p =* 0.002; Fig. S1).

We identified 127 volatile organic compounds (VOCs) in the communities across the gradient in plant species richness. These VOCs were classified into green leaf volatiles (GLVs) and other fatty acid derivatives, aromatics, nitrogen-containing compounds, homoterpenes, monoterpenes, sesquiterpenes, and a few other compounds that did not belong to any of these groups (Fig. 1). Sesquiterpenes and monoterpenes were the most numerous groups, with 41 and 28 compounds, respectively (Table S2). Since all plants have species-specific VOCs, we predicted that with increasing plant species richness the diversity of VOCs emitted from the communities will increase. Indeed, our results show that VOC emission (amount) and richness (number of compounds) increased with increasing plant species richness. However, this was not the case for VOC Shannon diversity or VOC Simpson diversity which suggests that while there are more compounds as plant species richness increases, their abundance does not become more even or dominant within the VOC profiles in the community (Fig. 2, emission: *x*^2^ = 7.84, *p =* 0.005; richness: *x*^2^ = 7.45, *p =* 0.006; Shannon diversity: *x*^2^ = 0.22, *p =* 0.64, Simpson diversity: *x*^2^ = 0.88, *p =* 0.35). Terpenoids were the main contributors to the increase in VOC emission (monoterpenes: *x*^2^ = 7.61, *p =* 0.006, sesquiterpenes: *x*^2^ = 6.35, *p =* 0.012, Fig. 2c-d), while the increase in VOC richness was mainly driven by sesquiterpenes (*x*^2^ = 7.46, *p =* 0.006, Table S3,4). Despite the fact that there was a strong positive correlation of plant phylogenetic diversity to species richness, VOC α diversity was not significantly influenced by phylogenetic diversity (Table S3).

**Figure 1.**
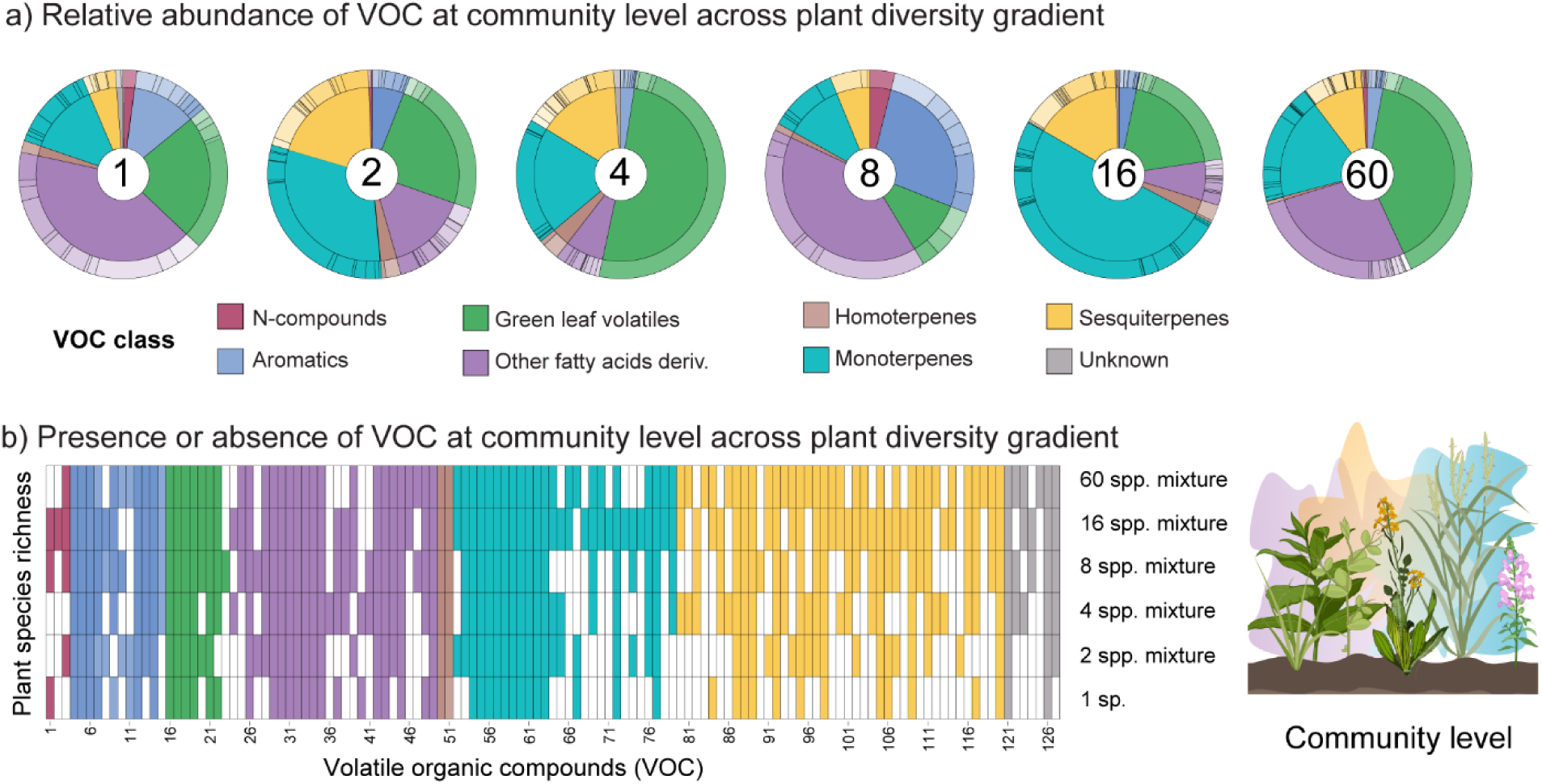
Volatile organic compound (VOC) profiles at the community level across the plant diversity gradient. Headspace VOC emission of experimental grassland communities was collected with a push-pull system (Figure S2) and analyzed by gas chromatography. a) Relative abundance of VOC classes across the experimental plant diversity gradient. Each pie chart depicts an example of one plant community from monoculture to 60 species mixtures. b) Presence (colored cell) or absence (white cell) of individual VOCs across the plant diversity gradient. Each column corresponds to a specific VOC compound, with compound names provided in Table S2.

**Figure 2.**
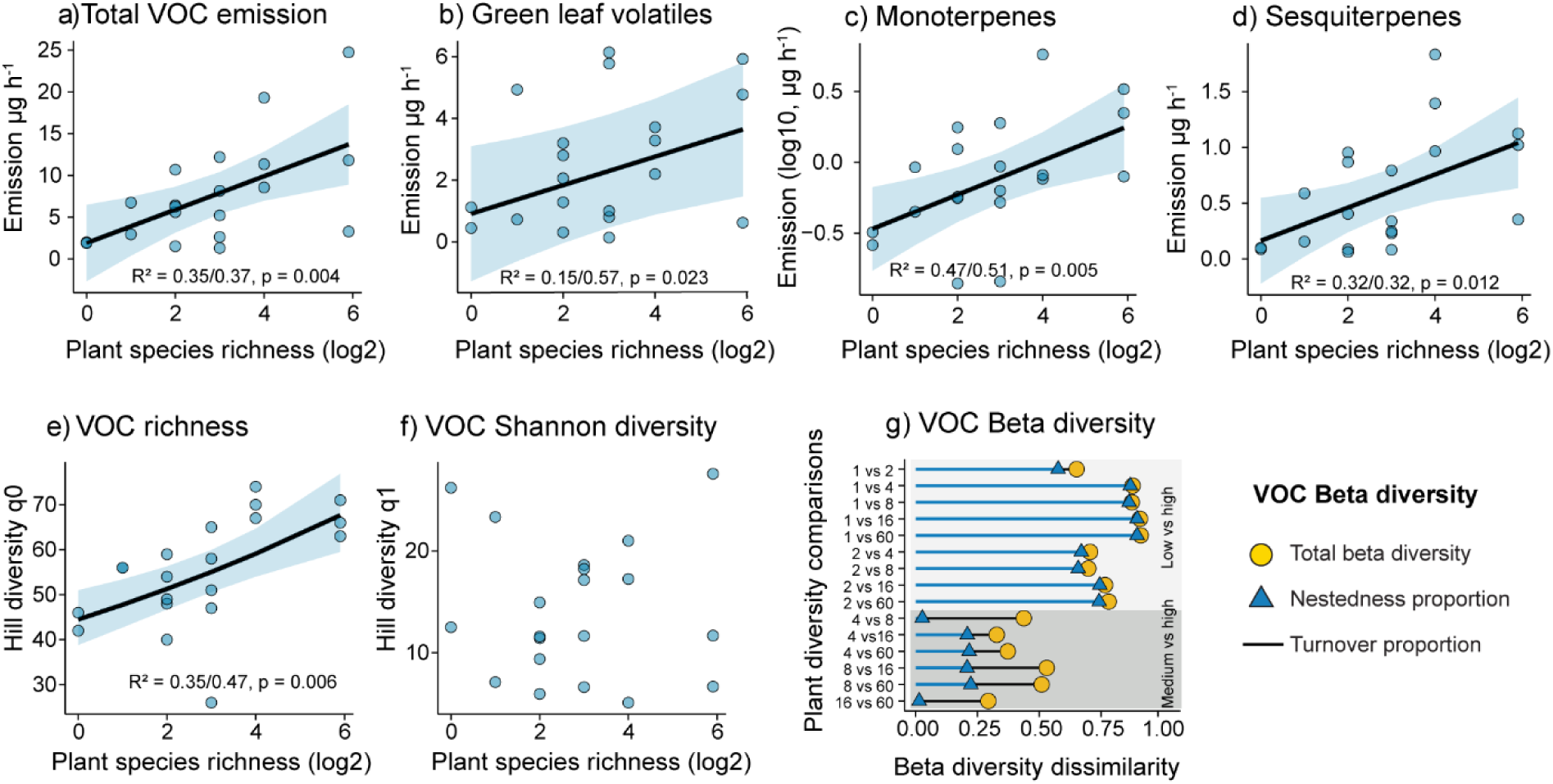
Volatile organic compound (VOC) emission and diversity at the community level across the plant diversity gradient. Headspace VOC emission of experimental grassland communities across a plant diversity gradient ranging from monocultures to 60-species mixtures. (a-d) VOC emission (µg per hour) at the community level, including a) total volatiles, b) green leaf volatiles, c) monoterpenes, and d) sesquiterpenes. (e-g) VOC diversity at the community level, including e) richness (Hill q0, number of compounds), f) Shannon diversity (Hill q1), and g) beta diversity across the diversity gradient. Sample size = 20 communities, with each data point representing the sum of VOC profiles from three random spots within each community. The lines show significant relationships (*p* < 0.05).

Since we observed an increase in the number of VOCs present in community VOC profiles as species richness increases, we used β diversity metrics to understand how VOC composition changes with the experimental plant diversity gradient. We found that VOC β diversity dissimilarity was higher when we compare low-diversity communities (1- and 2-species mixtures) to more diverse plant communities. This pattern was primarily due to nestedness, meaning that VOCs in low-diversity communities are mostly a subset of those found in more diverse communities. In contrast, among more diverse plant communities (4- to 60-species mixtures), the differences between communities were smaller. In these cases, we found that these communities had lower proportions of nestedness, and therefore VOC turnover better explained the VOC dissimilarities between communities. This implies a replacement of VOCs, which happens when the addition of new compounds balances out the absences of others (Fig. 2g). Most of the compounds that were involved in the high turnover proportion were sesquiterpenes.

To investigate the direct and indirect effects of biotic and abiotic conditions on community level VOC profiles across the diversity gradient, we incorporated data on Oomycota- and arbuscular mycorrhizal fungi (AMF) diversity, herbivore leaf damage, leaf area index, and soil temperature across the communities studied. We then constructed a Structural Equation Model (SEM; Fig. S2), which showed a good fit (Fischeŕs C = 20.33, p = 0.85, AIC = 524.33, Fig. 3) and explained a high proportion of the total variance in VOC emission and VOC richness (R_m/c_^2^ = 0.28 / 0.82 and R_m/c_^2^ = 0.64 / 0.64, respectively). A strong direct association between plant species richness and VOC profiles was identified even when accounting for the effects of biotic and abiotic factors. Phylogenetic plant diversity contributed again little to the VOC profiles but did promote Oomycota (soil pathogens) diversity, thereby marginally increasing VOC richness. Additionally, the LAI played a significant role in directly and indirectly shaping VOC richness in the community. Total VOC emission was mainly explained by plant species richness in the community and contributed marginally to a reduction in herbivore loads. Notably, increased plant species richness led to higher herbivore loads, which in turn promoted VOC richness. Soil temperature and arbuscular mycorrhizal fungi diversity did not significantly influence VOC profiles at the community level (Fig. 3, Table S5).

**Figure 3.**
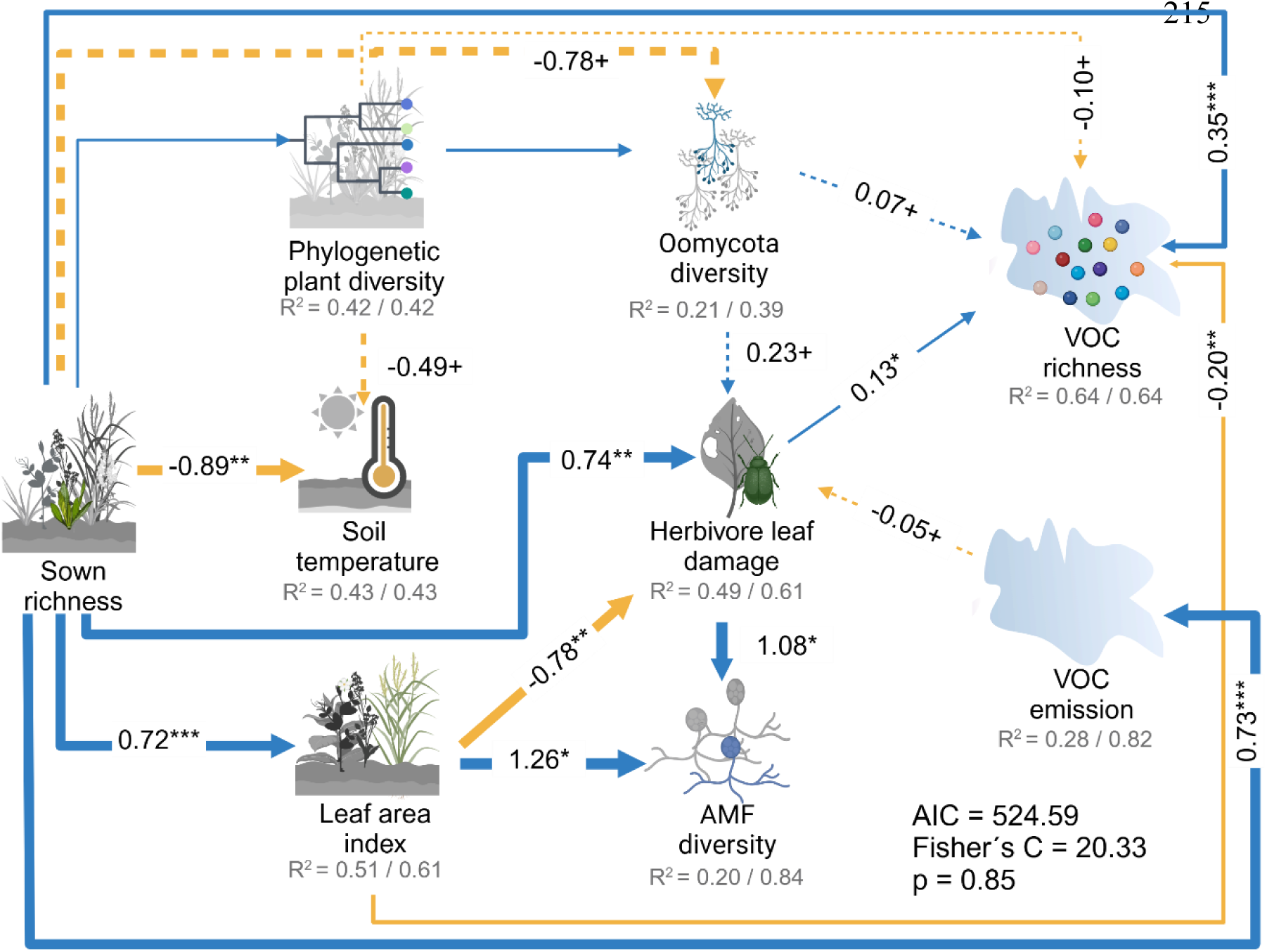
Results from the structural equation model (SEM) investigating the direct and indirect effects of biotic and abiotic conditions on VOC profiles at the community level across a diversity gradient. Blue arrows represent positive effects, while yellow arrows denote negative effects. The width of each arrow is proportional to the strength of the effects, as indicated by the standardized coefficient on each line. Only paths with statistical significance (*p < 0.05, **p < 0.01, ***p < 0.001, solid lines) or a tendency (+p < 0.1, dashed lines) are shown. Marginal and conditional R^2^ for component models are reported under the response variables (marginal before the slash and conditional after).

### Plantago lanceolata VOC profiles are shaped by the VOCs of the surrounding community

We next turned our attention to the pattern of VOC diversity at the level of a single species. From the 20 communities surveyed on the plant diversity gradient, we selected the ones where *P. lanceolata* is a resident species (eight communities) and collected VOCs in four *P. lanceolata* individuals per community. The actual plant species richness and phylogenetic diversity in the community were positively correlated with the originally sown plant species richness (richness: R^2^ = 0.68, *p =* 0.013; phylogenetic diversity: R^2^ = 0.90, *p <* 0.001).

We identified 29 VOCs emitted from *P. lanceolata*, classified into GLVs (5) and other fatty acids derivatives (4), aromatics (2), nitrogen-containing compounds (1), homoterpenes (1), monoterpenes (7) and sesquiterpenes (9) (Fig. 4, Table S2). Overall, the total emission and diversity of VOCs from *P. lanceolata* were not significantly influenced by the plant species richness of the surrounding community (Fig. 5a, e, I, Table S6). However, we found that the emission of the GLVs, (*Z*)-3-hexanol and (*Z*)-2-hexenyl acetate, and the monoterpenes, α-pinene and (*E*)*-*β-ocimene, decreased with the increase of plant species richness in the community (Table S7).

**Figure 4.**
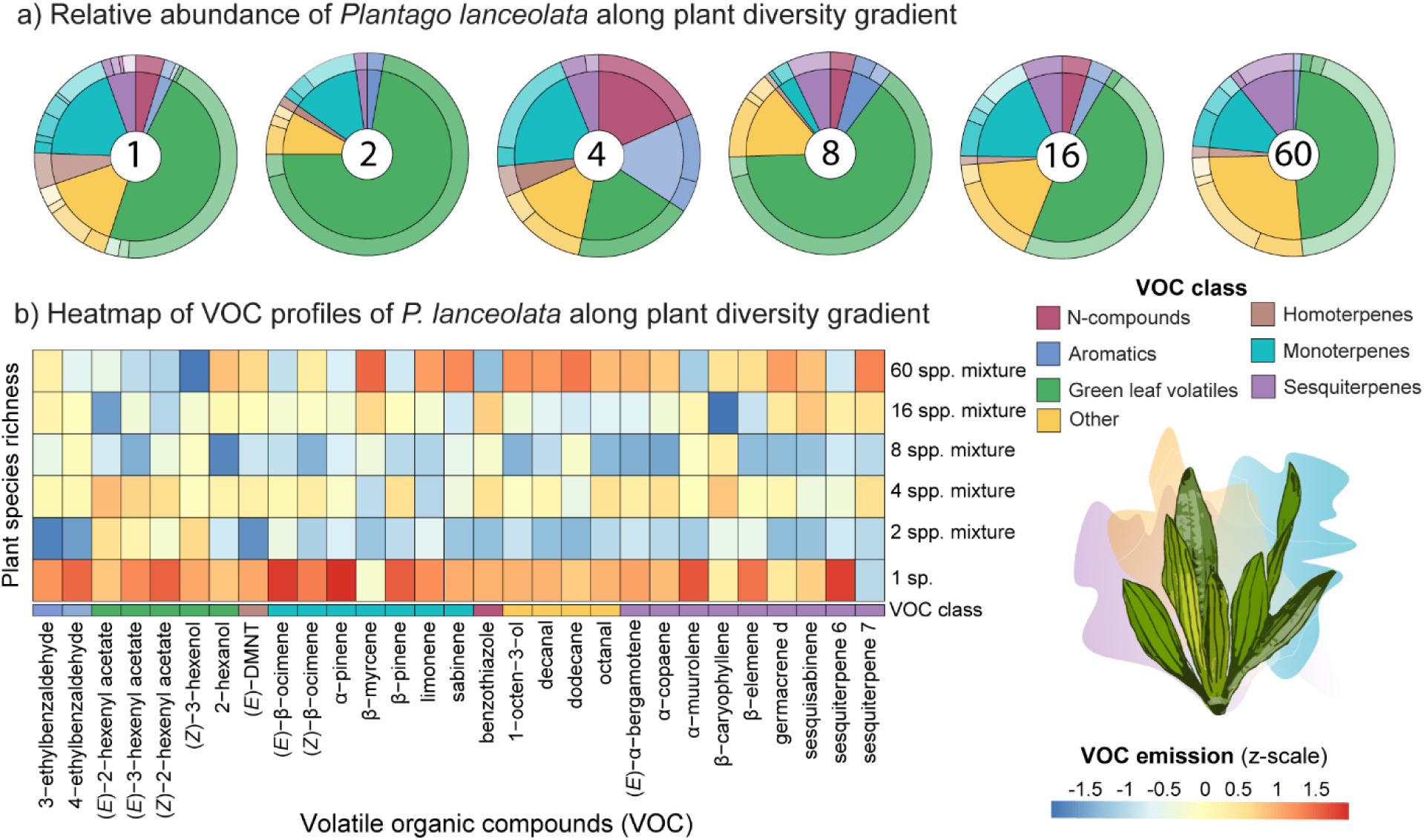
Volatile organic compound (VOC) profiles of *Plantago lanceolata* across the plant diversity gradient. a) Relative abundance of VOC classes emitted by *P. lanceolata* across a plant diversity gradient, spanning from *P. lanceolata* monoculture to 60-species mixtures. Each pie chart represents the relative abundance of different VOC classes of *P. lanceolata* in a community within the plant species richness gradient. b) A heatmap showing the relative mean abundances (z-scaled per compound across treatment) along the plant species richness gradient. Each column corresponds to a specific VOC compound.

**Figure 5.**
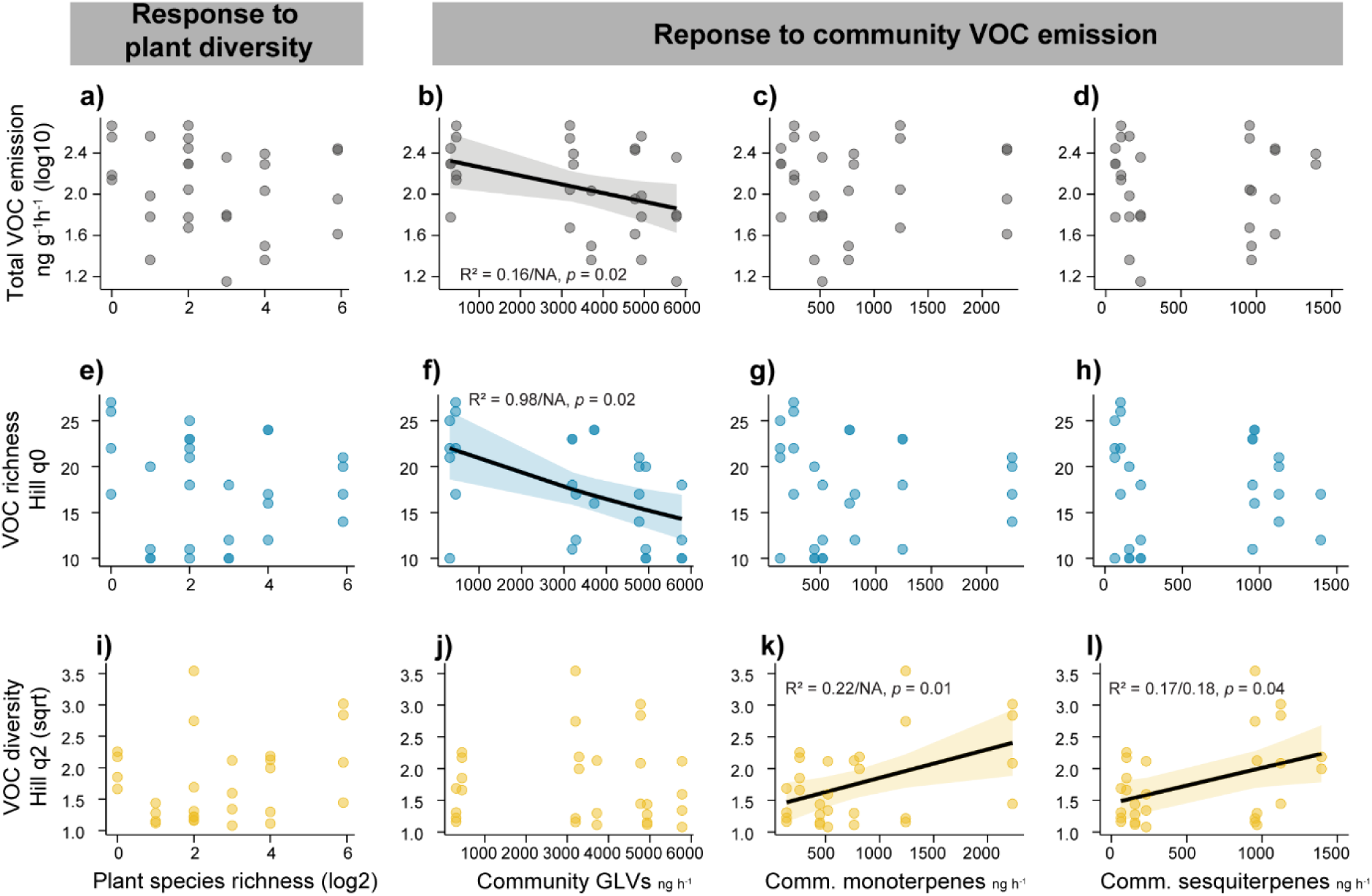
Volatile organic compound (VOC) emission and diversity of *Plantago lanceolata* in response to plant diversity and community VOC emission. Headspace VOC emission of resident individuals of *P. lanceolata. (a-d)* Total emission (grey, ng per g of fresh weight per hour) across (a) sown species richness, and the emission of (b) green leaf volatiles (GLVs), (c) monoterpenes and (d) sesquiterpenes in the surrounding community. (e-h) VOC richness (blue, Hill q0, number of compounds) across (e) sown species richness, and the emission of (f) GLVs, (g) monoterpenes and (h) sesquiterpenes in the surrounding community. (i-l) VOC diversity (yellow, Hill q2, Simpson diversity) across (i) sown species richness, and the emission of (j) GLVs, (k) monoterpenes and (l) sesquiterpenes in the surrounding community. The lines show significant relationships (*p* < 0.05). Sample size = 29 individuals.

When we investigated how the VOC profiles of the surrounding community influenced *P. lanceolata* VOC profiles, we observed notable effects. In general, an increase in GLV emissions in the surrounding community led to a decrease in both the total emission and number of VOCs emitted by *P. lanceolata* (Fig. 5b, f, Table S8). Specifically, *P. lanceolata* reduced the emission of GLVs, such as (*E*)-3-hexenyl acetate and (*Z*)-2-hexenyl acetate, and decreased the emission of terpenes such as, β-pinene, (*E*)-β-ocimene and α-muurolene; with a trend towards lower α-pinene, (*Z*)-β-ocimene and 4,8-dimethylnona-1,3,7-triene (DMNT) emission (Table S9). With increasing monoterpene and sesquiterpene emission in the surrounding community, the Simpson VOC diversity in *P. lanceolata* increased (Fig. 5k, l), which means that its VOC profiles became dominated by fewer compounds. This shift was primarily driven by the increased emission of (*E*)-2-hexenyl acetate, β-myrcene, α-muurolene and germacrene D, and the decreased emission of (*Z*)-3-hexenol and (*Z*)-2-hexenyl acetate (Table S9). Vegetation height of the surrounding community did not significantly influence *P. lanceolata* VOC emission or its diversity, but total GLV emission of *P. lanceolata* decreased with increasing vegetation height (*x*^2^ = 4.43, *p* = 0.035), specifically the emission of (*E* and *Z*)-2-hexenyl acetate.

Overall, total leaf damage in *P. lanceolata* decreased with increasing plant species richness (*x*^2^ = 5.37, *p* = 0.021; Table S6), driven primarily by a reduction in pathogen damage (*x*^2^ = 3.38, *p* = 0.06) rather than by a reduction in herbivore damage (*x*^2^ = 0.96, *p* = 0.33). When we compared the VOC profiles of *P. lanceolata* with the type of leaf damage, we found that VOC diversity was not influenced by herbivore leaf damage but rather by pathogen damage. VOC Simpson diversity decreased with increasing pathogen damage (*x*^2^ = 6.47, *p* = 0.011), especially due to a decrease in sesquiterpene emission (Table S10). Notably, herbivore damage decreased when individuals were surrounded by communities emitting a high concentration of GLVs (*x*^2^ = 4.19, *p* = 0.04; Table S8), while pathogen damage decreased when individuals were surrounded by communities emitting high terpene concentrations (monoterpenes*: x*^2^ = 7.36, *p* = 0.007; sesquiterpenes: *x*^2^ = 14.25, *p* < 0.001; Table S8).

## Discussion

In this study, we investigated how plant VOC emission varied across a plant diversity gradient, examining emission from both a whole plant community and from single individuals of *P. lanceolata.* Our results show that VOC diversity increased with increasing plant diversity at the community level, and the VOC emission of an individual plant species was affected more by the emission of the surrounding community than by plant species richness. Furthermore, at the species level, leaf pathogen damage decreased the VOC diversity of *P. lanceolata*. At the community level, plant species richness directly increased VOC emission, while its effect on VOC richness was mediated both directly and indirectly through changes in herbivore damage and LAI. The results show that the scale of observation is of crucial importance in determining which factors influence plant VOC diversity when a plant diversity gradient is altered.

Our results showed that high-diversity plant communities emit larger amounts, and a greater number of VOCs compared to low-diversity communities. This finding supports the hypothesis that introducing a new plant species into a community enhances the VOC profile by adding unique compounds, as VOC composition varies across different plant species^27,38^. Contrary to our expectations, increased plant phylogenetic diversity in the community did not significantly influence their VOC profiles. VOCs seem to transcend phylogenetic boundaries, as many compounds are shared across plant species regardless of their phylogenetic relationships^39^. However, while we found no significant relationship between plant phylogenetic diversity and VOC diversity, further studies at the community and species level are needed to explore the potential of phylogenetic diversity to alter the VOC profiles of plant communities.

Beta diversity analysis revealed that VOC profile dissimilarity increased with plant diversity in a community. As plant species richness increased, community-level VOC differences shifted from being primarily driven by compound nestedness (the presence or absence of compounds) to being driven by compound replacement (changes in the identity of emitted compounds). This suggests that there is a point where the total number of compounds stabilizes, but the identity of those compounds varies significantly. Interestingly, 37% of the VOCs were ubiquitous across all plant diversity levels studied, including compounds like (*Z*)-3-hexen-1-ol, α-pinene, (*E*)-β-caryophyllene, and methyl salicylate, which are often constitutively emitted or have their emission frequently induced by environmental stimuli from many plant species^40^. This variation in VOC composition driven by plant diversity may influence biotic interactions and structure in the community^41,42,^ emphasizing the potential role of VOC diversity in biodiversity-ecosystem functioning relationships. Assessing the ecological impacts of compound enrichment or replacement in VOC bouquets, however, can be a challenge, especially at the community level, as VOC functions may vary between species and organ type^43^. Furthermore, plants emit and perceive info-chemicals simultaneously, yet how these interactions change across different ecological scales remains largely unexplored, especially under field conditions.

Plants are known to respond to VOCs emitted by their neighbors^44^, and this response can be influenced by the identity of the neighboring plants and the diversity of the surrounding community^5,38,45^. In this study, plant diversity did not directly affect the VOC emission of resident *P. lanceolata* individuals, but plant diversity influenced the VOC emission of plant communities, which in turn influenced the VOC emission of *P. lanceolata* within them. Our findings add to previous studies on plant-plant communication^5,22,23,38,45^ by demonstrating that VOC emissions of receiver plants are influenced not only by individual neighbors but also by the overall VOC profile of the community. The observed decrease in total VOC emissions of *P. lanceolata* in communities with high concentrations of GLVs, together with their increased VOC diversity in communities dominated by terpenoids, may be attributed to several underlying explanations. For instance, 1) plants might prioritize alternative defensive strategies in "noisy" VOC environments, where signaling is less effective; 2) they might allocate fewer resources to VOC emission when benefitting from the protective effects of VOCs released by other plants in the community; 3) the emission of fewer VOCs may reduce the risk of eavesdropping by neighboring plants, thereby enhancing the defensive advantage of the emitter; or 4) these changes could represent adaptive responses to specific VOC cues conveying different kinds of information. Since the diversity of terpene volatiles emitted by plants is often far greater than the diversity of GLVs, terpenoids may transmit more distinct information^43,46,47^. Certainly, further research is needed to fully understand the basic functions of plant-plant communication and how this can vary across diversity gradients.

In this study, *P. lanceolata* individuals experienced lower herbivore damage in communities emitting high concentration of GLVs, and pathogen damage decreased when individuals were surrounded by communities emitting high terpene concentrations. These results suggest that plants exhibit specific responses to distinct VOCs in the community^48^. The surrounding VOC composition can influence *P. lanceolata* performance by direct effects or by modifying plant defense strategies. For instance, exposure to GLVs and terpenes from neighboring plants not only alters the VOC emission patterns of receiver plants^49^ but also induces the accumulation and biosynthesis of defensive compounds, reducing herbivore and pathogen damage^50–52^. Moreover, at the community level, VOCs function as long-distance signals, attracting or repelling organisms attempting to enter the community, while either facilitating or disrupting communication among plant species^45^. Increased VOC emissions and diversity may disrupt cues used by specialist herbivores or pathogens, benefitting plants through "chemical noise"^53^, but greater VOC complexity might also hinder interactions with mutualists like pollinators or parasitoid wasps^10,54^. Surrounding VOCs could also have direct effects on pathogens or herbivores in sufficient concentrations since many of these compounds are known to be toxins or feeding deterrents^55^. Overall, VOC profiles might play a crucial role in shaping biotic interactions, directly enhancing plant defenses and indirectly influencing ecological dynamics.

As mentioned before, both antagonistic and mutualistic organisms can shape plant VOC emissions^9,56,57^. More importantly, in diverse plant communities, there is often an accumulation of mutualists and a dilution of antagonists, both above and below ground^28,58,59^. These shifts can drive individual plants to emit distinct VOC profiles in low-*versus* high-diversity communities due to varying pressures^5^. However, most existing studies have focused on species -level interactions, and scaling up plant VOC-mediated interactions to the community level can be a challenge. Increasing chemical diversity in the community can be positively correlated with the abundance of arthropods and reduce leaf damage.^60^. In this study, an increase in plant species richness shaped VOC profiles in the community through both direct and indirect pathways. While VOC emission was directly enhanced by higher plant species richness rather than biotic factors, VOC richness was influenced not only directly by plant species richness but also indirectly through its effects on LAI, herbivore damage, and soil microbiota. Thus, different community traits shaped VOC differently, with plant richness directly enhancing VOC emission, while its influence on VOC richness also occurs through modifications in their abiotic and biotic factors. Given the fact that many VOCs produced by plants can also be emitted by other organisms such as microbes or insects^61,62,^ we should be aware that the VOC bouquet we sampled may not be exclusively plant-derived.

In conclusion, our findings provide first insights into how plant diversity shapes VOC emissions at both the species and community level. Higher plant species richness within a community increases the number and diversity of VOCs emitted, shifting from richness-driven changes to compound-specific replacements as plant diversity increases. Additionally, we also showed that community-level VOCs influence individual plant emissions and leaf damage. Our results highlight the critical role of VOC diversity in plant-plant communication and interactions with higher trophic levels, emphasizing its potential impact on biodiversity and ecosystem functioning. Further research in this field is needed to understand the underlying mechanisms by which community-level VOCs could influence plant performance by direct or receiver plant-mediated interactions.

## Material and methods

### Field site and experimental design

This study was conducted in the “The Jena-Experiment” (50°55’ N, 11°35’ E; 130 m a. s. l.), a long-term grassland biodiversity experiment in Jena, Germany^35^. Since 2002, the plant species community compositions, selected from a pool of 60 native grassland species, have been maintained through regular weeding (2–3 times per year) and mowing (twice yearly) to replicate traditional management of Central European grasslands. We selected 20 communities (plots) representing a gradient in species richness, ranging from monocultures to 60 plant species-mixtures (1, 2, 4, 8, 16, and 60 plant species), as well as different functional group richness (1 to 4 functional groups: grasses, small herbs, tall herbs, and legumes). Among the 20 communities studied, *Plantago lanceolata* L. (ribwort plantain) belonged to the sown species combinations in eight communities. At the community level, we measured VOC emission, realized plant species richness, aboveground biomass, vegetation height, leaf area index, soil temperature, herbivore damage and soil microbiota (oomycota and arbuscular mycorrhiza fungi). At the species level (*P. lanceolata*), we measured VOC emission, aboveground biomass and percentage of leaf damage by herbivores and pathogens.

### Headspace VOC collection

Volatile organic compound (VOC) emission at the community and species level was measured using a push-pull system for 2 hours (Figure S2). At the community level, three cylindrical frames (Ø 50 cm x 80 cm) covered with PET films (sealed with a film sealing machine FERNANT® 120 N) were installed randomly in the core area (3 x 3 m) of each community. At the species level, four individuals of *P. lanceolata* were individually enclosed with PET bags (Toppits® Bratschlauch, Melitta, Minden, Germany) tightened on top and at the bottom with a cable binder. A small sponge was used between the bag and the cable binder, to reduce potential damage of the plant. In both systems, charcoal-filtered air was continuously pumped into these cages or bags (Community level: 2 L/min flow rate, species level: 1 L/min). At the same time, air was pumped out through VOC traps, consisting of 25 mg of Porapak absorbent (ARS, Grainville, FL, USA) inserted in Teflon tubes (Community level: 1.0 L/min flow rate, species level: 0.7 L/min). All volatile collections were performed between 9:00 am and 1:00 pm. After VOC collection, the VOC traps were eluted with 200 µl dichloromethane containing nonyl acetate as an internal standard (SigmaAldrich, 10 ng µl^-1^). VOCs were analyzed using a gas chromatograph coupled to a mass spectrometer (GC-MS) with helium as the carrier gas for compound identification (Hewlett-Packard 6890 gas chromatograph coupled with a Hewlett-Packard 5973 mass spectrometer) and a gas chromatograph coupled to a flame ionization (GC-FID) detector with hydrogen as the carrier gas for compound quantification (details in ^63^). VOCs were identified by comparing retention times and mass spectra to those of authentic standards obtained from Fluka (Seelze, Germany), Roth (Karlsruhe, Germany), Sigma (St. Louis, MO, USA) or Bedoukian (Danbury, CT, USA) or to reference spectra in the Wiley and National Institute of Standards and Technology libraries (NIST). The quantity of each compound was determined from its peak area in the FID trace in relation to the area of the internal standard using the effective carbon number concept^64^.

### Community plant trait measures

In each community, we measured species-specific plant cover (% per species), plant aboveground biomass inside the VOC cages (fresh weight), plant community mean height and leaf area index (LAI). Percentage cover per species was estimated in a 3m x 3m area of the community using a decimal scale^65^, including all target species (sown in a particular community) and all plant species, which colonized the communities from the surroundings. Community leaf area index (LAI) was measured using a portable LAI-2200C plant canopy analyzer (LI-COR, Lincoln, USA) at ten randomly selected positions within each plot. Community mean height was calculated by averaging the vegetation height (highest leaves) measured at ten randomly selected spots in each plot. Based on the realized pant species richness and vegetation cover, we calculate plant taxonomic, functional, and phylogenetic diversity.

### Community herbivore damage

Herbivory damage by invertebrates and small mammals was quantified as the proportion of damaged leaf area relative to the total leaf area for each plant species within the community. To estimate community-level herbivory, a weighted mean of species-specific herbivory values was calculated for each plot, using leaf biomass as the weighting factor (details in ^58^).

### Community soil microbiota diversity

Soil Oomycota and arbuscular mycorrhizal fungi diversity were estimated at the community level from bulk soil collected in each studied community during the same period of VOC collection. Oomycota and AMF diversity were obtained from Albracht et. al^28^ and Li et al.^66^, and then transformed to Hill numbers (Hill q1). Briefly, DNA was extracted from bulk soil (pooled and homogenized from 4 random soil cores (4 cm diameter, 5 cm depth) per community) with the Quick-DNA Fecal/Soil Microbe Miniprep Kit (Zymo Research Europe GmbH, Freiburg, Germany). The ITS1 region of Oomycota was then amplified with barcoded primers applying the procedure described in Fiore-Donno and Bonkowski^67^ Sequencing was performed with a MiSeq v3 Reagent kit of 600 cycles on a MiSeq Desktop Sequencer (Illumina Inc., San Diego, CA, USA) at the Cologne Center for Genomics (CCG, Cologne, Germany). The raw sequence reads were assembled and quality-filtered in *mothur*^68^. Oomycota diversity (Shannon diversity) was estimated with *vegan*^69^. For AMF, the SSU rRNA region was amplified with nested PCR (primer pairs WT0/Glomer1536 and NS31 / AML2) and sequenced at the Illumina MiSeq platform at the Soil Ecology department of Helmholtz Centre for Environmental Research (Halle (Saale), Germany). Sequencing data was processed with the dadasnake pipeline^70^. AMF diversity (Shannon diversity) was estimated with *phyloseq*^71^ on rarefied data (2400 reads per sample).

### Measurement of soil temperature

In each plot, the soil temperature was measured at 5 cm, using thermometers of a controller area network-bus module system (JUMO). The temperature sensors were lance probes with a diameter of 4.5 mm and a length of 200 mm. The measuring element was a PT100 resistor with a tolerance of ± 0.1 °C at 0 °C. The sensor operated in a four-wire connection to the data-acquisition module of the controller area network-bus network^72^.

## Data analysis

Realized plant diversity was assessed at both the taxonomic and phylogenetic levels using Hill numbers. Plant species-level cover was used as an abundance variable for diversity calculations. Phylogenetic diversity was calculated by constructing a phylogenetic tree that included all plant species recorded across communities. To examine whether sown species richness was still significantly correlated to the realized plant diversity in the selected communities, we performed a linear regression model with sown species richness (log2 transformed) as the explanatory variable and the different plant diversity indices measured as dependent variables.

To explore the role of plant diversity in shaping volatile organic compound (VOC) emissions, we calculated VOC diversity at both the community and species levels. The alpha diversity metric was used to determine the relationship between the number of VOC compounds in a bouquet and their relative abundance, whereas the beta diversity metric was used to determine how VOC composition changes across the plant diversity gradient and whether this is due to compound turnover among communities or nestedness. VOC emission area was used as abundance information. Since the plant communities have different degrees of homogeneity, we summarize the emission of the three replicates of each community (ng h^-1^) and calculate their VOC diversity. At the species level, VOC diversity was standardized by leaf biomass to account for variation in plant size (ng g_fw_ h^-1^).

To evaluate the effects of plant diversity on VOC emission and diversity at the species- and community levels, we performed mixed-effect models. At the community level, we included fresh weight biomass as a covariable fitted before the fixed factor (sown plant richness or realized diversity), and block and collection date as random effects (e.g., y ∼ biomass + log2(sown plant richness) + (1|block) + (1|collection_date)). At the species level, we fitted either sown species richness, community VOC profiles, or leaf damage as fixed factors and plot nested in block and collection date as random effects (e.g., y ∼ log2 (sown plant richness) + (1|block/plot) + (1|collection_date)). To assess term significance, we used ANOVA type I sum-of-squares and adjusted p-values for false discovery rate (FDR) due to multiple testing^73^. We extracted marginal and conditional R² values from the models to evaluate the proportion of variance explained. When needed, data was transformed to meet the assumptions of normality.

To evaluate whether abiotic and biotic interactions driven by species diversity altered the effects and direction of the relationships on VOC emission and richness, we constructed a structural equation model (SEM) fitting mixed-effect models, following the conceptual framework described in Fig. S2. We selected the final model by excluding the non-significant factors and adding missing relationships that significantly improved the model’s global fit and were ecologically meaningful.

All statistical analyses and data visualization were conducted in R version 4.3.3^74^ using the *rBExIS, dplyr*, *tidyverse*, *tibble* and *janitor* for data retrieval, cleaning and formatting^75–78^. Plant diversity calculations were performed using the *hillR* package^79^, and phylogenetic diversity was estimated using the *V.PhyloMaker* package^80^, which constructs phylogenies based on the GBOTB backbone^81^. VOC diversity metrics were computed using the *hillR* and *BAT* packages^79,82^. Mixed-effects regression models were implemented using the *lme4*, *lmerTest* and *glmmTMB* packages^83–85^, with model variance components extracted using *MuMIn*^86^. Structural equation modeling was conducted using the *piecewiseSEM* package^87^. For data visualization, we used *ggplot2*, *ggeffects*, *webr*, and *pheatmap*^88–91^. Illustrations were created in R and BioRender.com, with final graphics refined in Adobe Illustrator CC 2021.

## Author’s contributions

P.M.B., and S.B.U. designed the study; P.M.B. performed the experiment, chemical analysis, and data analysis; A.H.B., M.B., F.B., C.A. and M.D.S. designed and carried out the Oomycota and mycorrhiza profiling at the community. S.M., A.E. and M.B. designed and carried out the herbivore damage sampling at the community level. A.E. and N.E. contributed vegetation cover and leaf area index data. K.K., T.E. installed and maintained the soil temperature measurement system. G.S., Y.H., cleaned and analyzed the soil data. N.E. and A.E. coordinate the Research Unit of the Jena Experiment. P.M.B. wrote the first draft of the manuscript. S.B.U. and J.G. revised the manuscript, and all co-authors discussed the results, contributed substantially to the drafts, and gave final approval of the manuscript prior to the submission.

## Acknowledgements

We are grateful to Nils Gottschaldt, Klara Beier-Heuchert, Melanie Werlich, Christiana Voy and Thimea Truepschuch for their help with the field experiments. We thank Daniel Veit for designing and building the volatile collection system. We also thank Annette Jesch for coordinating central data collection and weeding of the experimental plots, and the gardeners and student helpers for their valuable efforts in maintaining the field site. We also thank Christiane Roscher for her helpful comments on an earlier version of the manuscript. This study was funded by the German Research Foundation (FOR5000, UN276/4-1, EB555/6-1, ME5474/1-1). Moreover, N.E. and Y.H. acknowledge funding by the German Research Foundation (DFG, FZT 118, 202548816; Ei 862/29-1).

## Data availability

The data and R code will be publicly available through the Jena Experiment database (https://jexis.idiv.de). Datasets at the community level, including plot information (ID = 90), realized plant species at the plot level (vegetation cover; ID = 264) and inside the VOC cages (fresh weight biomass; ID = 682), mean height (ID = 346), leaf area index (ID = 290), soil temperature (ID = 326), soil pathogen diversity (oomycota; ID = 538), arbuscular mycorrhizal fungi diversity (ID = 319), percentage of leaf damage by herbivores (ID = 364), and VOC profiles (ID = 726) will be publicly available upon acceptance. At the species level, VOC profiles (ID = 725) and percentages of leaf damage by herbivores and pathogens (ID = 685) will be publicly available upon acceptance. Upon acceptance, we will deposit the R codes used in BEXIS.

## Supplemental information

- **Figure S1.** Realized plant species diversity of the communities studied across a diversity gradient in the Jena Experiment during the summer of 2021.
- **Figure S2.** Push-pull systems for volatile organic compounds collection in the field.
- **Figure S3.** Conceptual framework of the direct and indirect effects of plant species diversity in plant volatile organic compound emission in an experimental grassland field.
- **Table S1.** Plant species in the communities studied of the Jena Experiment in May 2021.
- **Table S2.** List of volatile compounds identified at plant community level and species level.
- **Table S3.** Community level: Wald-chi-squared analysis of variance (ANOVA) results for the linear mixed-effects models testing effects of plant diversity on VOC emission and diversity at community level.
- **Table S4.** Community level: Wald-chi-squared analysis of variance (ANOVA) results for the linear mixed models testing effects of plant diversity on each VOC emitted at community level.
- **Table S5.** Community level: Path coefficient for the fitted final structural equation model to investigate the direct and indirect effects of plant species richness on plant volatile profiles in an experimental grassland field.
- **Table S6.** Species level: Wald-chi-squared analysis of variance (ANOVA) results for the linear mixed models testing the effects of plant diversity on *Plantago lanceolata* VOC emission and leaf damage.
- **Table S7.** Species level: Wald-chi-squared analysis of variance (ANOVA) results for the linear mixed models testing the effects of plant diversity and leaf damage on each VOC emission of *Plantago lanceolata*.
- **Table S8.** Species level: Wald-chi-squared analysis of variance (ANOVA) results for the linear mixed models testing the effects of Community VOC on *Plantago lanceolata* VOC emission and leaf damage.
- **Table S9.** Species level: Wald-chi-squared analysis of variance (ANOVA) results for the linear mixed models testing the effects of Community VOC on each VOC emission of *Plantago lanceolata*.
- **Table S10.** Species level: Wald-chi-squared analysis of variance (ANOVA) results for the linear mixed models testing the effects of leaf damage on *Plantago lanceolata* VOC emission.

